# Differential Patterns of Associations within Audiovisual Integration Networks in Children with ADHD

**DOI:** 10.1101/2023.09.26.559610

**Authors:** Mohammad Zamanzadeh, Abbas Pourhedayat, Fatemeh Bakouie, Fatemeh Hadaeghi

## Abstract

Attention deficit hyperactivity disorder (ADHD) is a neurodevelopmental condition characterized by symptoms of inattention and impulsivity and has been linked to disruptions in functional brain connectivity and structural alterations in large-scale brain networks. While anomalies in sensory pathways have also been implicated in the pathogenesis of ADHD, exploration of sensory integration regions remains limited. In this study, we adopted an exploratory approach to investigate the connectivity profile of auditory-visual integration networks (AVIN) in children with ADHD and neurotypical controls, utilizing the ADHD-200 rs-fMRI dataset. In addition to network-based statistics (NBS) analysis, we expanded our exploration by extracting a diverse range of graph theoretical features. These features served as the foundation for our application of machine learning (ML) techniques, aiming to discern distinguishing patterns between the control group and children with ADHD. Given the significant class imbalance in the dataset, ensemble learning models like balanced random forest (BRF), XGBoost, and EasyEnsemble classifier (EEC) were employed, designed to cope with unbalanced class observations. Our findings revealed significant AVIN differences between ADHD individuals and neurotypical controls, enabling automated diagnosis with moderate accuracy. Notably, the XGBoost model demonstrated balanced sensitivity and specificity metrics, critical for diagnostic applications, providing valuable insights for potential clinical use.

These findings offer further insights into ADHD’s neural underpinnings and highlight the potential diagnostic utility of AVIN measures, but the exploratory nature of the study underscores the need for future research to confirm and refine these findings with specific hypotheses and rigorous statistical controls.

## 1 Introduction

Attention Deficit Hyperactivity Disorder (ADHD) is associated with sensory processing deficits, including difficulties in modulating sensory information across various domains [1, 2]. Notably, children with ADHD often display heightened distractibility, particularly in response to auditory stimuli in audiovisual (AV) conditions, which reflects a challenge in inhibiting irrelevant inputs [3, 2].

While previous research has highlighted sensory processing deficits in ADHD, the investigation of audiovisual integration (AVI) in children with this condition remains limited. Studies have primarily focused on AVI in adults with ADHD, indicating impairments in multisensory processing due to underlying unisensory deficits [4, 5, 6]. However, the understanding of AVI in the context of ADHD specifically in children is still in its infancy.

One possible reason for this research gap is the late development of multisensory integration abilities. Generally, neurotypical children begin to exhibit multisensory integration after the age of four, with optimal integration not fully developed until late childhood, around ages 8 to 10 years [7]. This suggests that children with ADHD might also experience delayed or impaired AVI compared to their neurotypical peers.

Understanding AVI deficits in children with ADHD is further complicated by the lack of consensus regarding the specific brain regions involved in multisensory integration. While certain regions, such as the superior temporal sulcus (STS) [8, 9], superior parietal sulcus (SPS) [10, 11], intraparietal sulcus (IPS) [12], inferior parietal gyrus (IPG) [12, 11], insula (INS) [11], Heschl’s gyrus (HES), and calcarine fissure and its surrounding cortex (CAL) [13], have been implicated in AVI during task performance, ongoing research is needed to establish a comprehensive understanding of the neural mechanisms underlying AVI.

To address these research gaps, we conducted a novel investigation utilizing resting-state functional magnetic resonance imaging (rs-fMRI) data and graph theory-based analyses. Our study aimed to explore network connectivity patterns and extract relevant features within the AVI-related brain network in children with ADHD (average age: 9.5 years). The primary objective was to identify potential diagnostic biomarkers for ADHD based on AVI, thereby enhancing our understanding of sensory processing differences in this population.

First, we identified a network of brain regions known to be associated with AVI, including the superior temporal gyrus/sulcus (STG), superior parietal sulcus (SPS), Heschl’s gyrus (HES), and calcarine fissure and its surrounding cortex (CAL). We selected these regions based on their potential role in AVI during resting state, as supported by previous literature [8, 9, 10, 11, 13, 12].

Next, we extracted graph theoretical-based features from the connectivity patterns within this network. These features provided quantitative metrics capturing the organization and efficiency of the AVI-related brain network in each participant. To explore group differences, we compared the extracted features between a control group and children with ADHD using classical statistical analyses. This allowed us to identify significant statistical differences that could highlight distinctive patterns of AVI-related brain connectivity in children with ADHD.

In addition to the graph theoretical analysis, we employed network-based statistics (NBS) as a complementary approach. Our NBS analyses focused on the raw connectivity matrices to identify a sub-network composed of specific nodes and their pairwise connections exhibiting significant differences between the two groups.

Importantly, we went beyond the descriptive and statistical analyses by applying machine learning techniques. Instead of using the full-resolution 3D spatial structure of rs-fMRI to deep learning models [14], we used the extracted graph features as inputs and trained classification models to distinguish between the control group and children with ADHD. We evaluated the diagnostic potential of AVI-related rs-fMRI data and graph theory-based features as potential biomarkers for ADHD. This involved splitting the data into training and validation sets, iteratively training the models on different subsets of the data, and evaluating their performance on the validation sets. Additionally, we used a held-out test set that was not used during the cross-validation process to assess the generalization ability of the selected model. Notably, when comparing the performance of these models on the default mode network (DMN), which is commonly studied in ADHD, we found that our classifiers trained on the AVI network outperformed them. This indicates the potential superiority of AVI-related features in capturing distinctive patterns of ADHD.

By combining rs-fMRI data, graph theory, statistical analyses, and machine learning, our study aimed to provide a comprehensive understanding of AVI in children with ADHD. We investigated both the group differences in AVI-related brain connectivity and the potential of using these connectivity patterns as a diagnostic tool. The findings from our study could significantly contribute to the development of objective and quantitative biomarkers for ADHD, offering valuable insights for diagnosis and potential interventions.

## 2 Methods

In this section, we elucidate the procedures and techniques, as exemplified in Figure 1, applied in this study.

**Figure 1:**
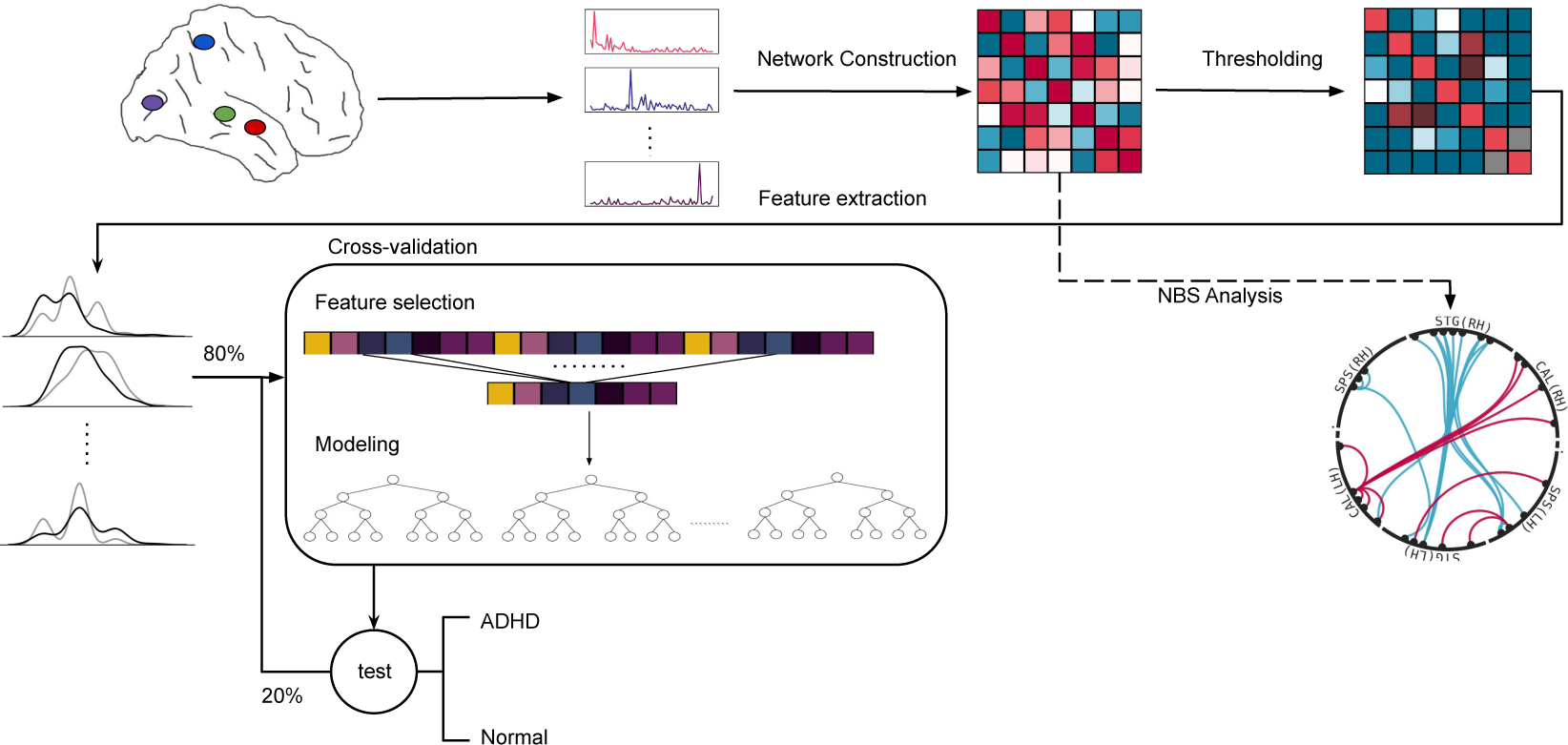
Schematic representation of our pre-processing and data analysis pipeline.

### 2.1 Dataset

We conducted our study on the ADHD-200 Preprocessed dataset [15], which was made available as part of the ADHD-200 global competition and since then has been extensively used in childhood ADHD research. This dataset comprises a total of 973 participants who were scanned at eight different sites. Among these participants, there were 585 individuals classified as neurotypical, 362 individuals diagnosed with ADHD and 26 participants whose diagnosis was unavailable.

To ensure data collection consistency, participants at each site followed specific instructions to either keep their eyes closed or open during the scanning procedure [15]. Notably, the Kennedy Krieger Institute, Brown University, and Oregon Health and Science University sites had explicit guidelines requiring all participants to keep their eyes open or fixated. Considering our focus on investigating audio-visual integration (AVI), we selected data exclusively from the Kennedy Krieger Institute and the Oregon Health and Science University sites. Unfortunately, due to unavailable true labels of participants from the Brown University site, the data collected from this center could not be utilized in our study.

After excluding left-handed individuals, we further refined the dataset to only include participants at an appropriate developmental stage, resulting in a final sample size of 180 participants. Among them, there were 127 neurotypical individuals (64 females) with an average age of 9.80*±*1.38 years. The remaining 53 participants were diagnosed with ADHD (36 females) and had a mean age of 9.59 *±* 1.44 years.

The rs-fMRI data has already been preprocessed using the pipeline available as part of the Neuroimaging Analysis Kit (NIAK) for Octave and Matlab [16]. This pipeline encompasses several essential steps, including the removal of the initial four volumes, correction of slice timing, realignment of volumes with respect to the first volume, registration of rs-fMRI and s-MRI using a linear transformation, application of a bandpass filter (with cutoff frequencies of 0.0009 Hz and 0.08 Hz), and spatial smoothing through a 6.0 *mm* full-width at half-maximum (FWHM) Gaussian filter. These procedures aim to eliminate artifacts, correct for temporal discrepancies, enhance signal quality, and reduce noise originating from low and high frequencies within the data.

### 2.2 Regions of interest

The preprocessing pipeline employs a spatially constrained spectral clustering to generate a whole brain fMRI atlas consisting of 954 functional regions of interest (ROIs) [17]. In our study, we selected the 68 ROIs from both hemispheres for investigating the audiovisual integration (AVI) network based on the anatomical regions defined by the automated anatomical atlas 3 (AAL3) [18]. The AAL3 atlas provides detailed anatomical parcellations of the brain, and we chose regions that are relevant to audio-visual integration. Specifically, we focused on regions that are known to be involved in multisensory processing according to their anatomical definitions in the AAL3 atlas. Additionally, to ensure a sufficient signal-to-noise ratio in our functional connectivity analysis, we considered regions with at least 100 voxel size. This criterion helped us select regions that are robust and well-defined for our study’s objectives. The specific coordinates of these ROIs can be found in Supplementary Table S2 for reference.

The selected ROIs, depicted in Figure 2, were distributed across four distinct areas of the brain: the superior temporal gyrus (STG), the Heschl’s gyrus (HES), the calcarine sulcus (CAL), and the superior parietal sulcus (SPS). In the context of rs-fMRI, we deliberately excluded brain regions that exhibit activation exclusively during audio-visual sensory integration tasks. Instead, the focus has been directed toward brain regions that are hypothesized to engage in audio-visual integration even during periods of rest. The designated cortical areas under investigation are those that demonstrate heightened activity during the concurrent and spatially co-localized processing of both visual and auditory stimuli. Notably, their function pertains to the examination of the inherent attributes intrinsic to the presented stimulus, rather than being centered on its content-related characteristics.

**Figure 2:**
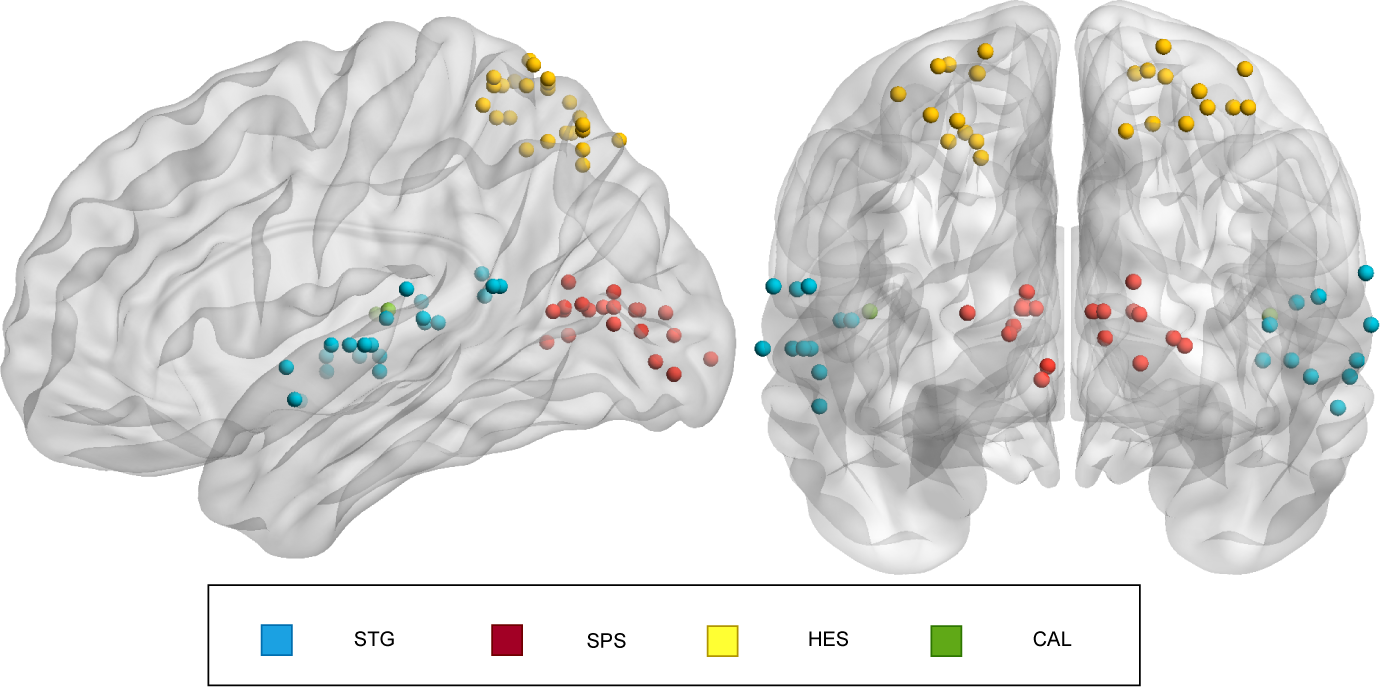
Sagital (left) and coronal (right) view of the brain. Different ROIs are shown. We employed the MNI atlas to define regions of interest. Brainnet viewer software package (http://nitrc.org/projects/bnv/) was used to draw this picture.

To perform further analysis, regional time series were extracted by averaging the signal obtained from the voxels within each of these regions. These regional time series were then utilized for subsequent investigation and analysis.

### 2.3 Connectivity matrix construction

We explored two different methods to measure the interactions between regional signals. The first method involved linear Pearson correlation, where the weighted functional connectivity (*W_ij_*) between two nodes (*i* and *j*) is determined by calculating the Pearson correlation coefficient between their respective time courses throughout the entire scanning duration. The second method involved using non-linear mutual information (MI), which assesses the interdependence between two random variables and measures how much knowledge about one variable reduces the uncertainty about the other variable [19]. To compute mutual information, we employed the following formula that quantifies the interdependence between two random variables, *X* and *Y* :

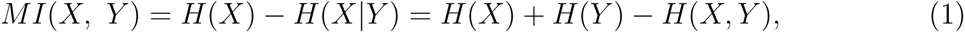

where *H*(*X*) denotes Shannon entropy of the variable *X*. Unlike Pearson correlation, mutual information takes into account both linear and nonlinear relationships and does not yield negative values.

To suppress spurious edges in the reconstructed networks, we employed a density-based thresholding method that preserved a specified percentage, denoted as *τ*, of the total edges, serving as a flexible parameter in our study.

Our analysis commences with a rigorous examination of functional connectivity profiles through the application of the network-based statistics (NBS) test, as outlined by Zalesky et al. [20]. In the context of case-control studies, the NBS method serves the purpose of identifying a sub-network composed of nodes and their pairwise associations within connectivity patterns that exhibit notable disparities between groups. Each association is assigned a test statistic, and in certain cases, a transformation may be employed to refine the raw measure of association. The central focus of the NBS approach lies in the identification of interconnected structures, accomplished by selecting suprathreshold links and assessing their topological scope to ascertain their significance.

To determine the statistical significance of these identified structures, permutation test-ing is employed. This entails generating numerous random permutations by interchanging the group assignments of the subjects, followed by recalculating the test statistics for each permutation. A threshold is then applied to establish a set of suprathreshold links for every permutation. Within each permutation, the maximum component size among the suprathreshold links is determined and stored, allowing for the creation of an empirical estimation of the null distribution. By comparing the observed component size against this null distribution, a *p*-value is computed to assess the significance of the identified component. This permutation testing approach, seamlessly adapted to the graph model, offers an effective means of identifying and evaluating interconnected structures within the framework of case-control studies.

### 2.4 Feature extraction

In our analysis of AVI networks, we further employed a diverse range of nodal and network-wide graph theoretical measures on the reconstructed connection matrices. These measures encompassed various classes, including node degree, centrality, assortativity, modularity, communication and information flow efficiency, and more, offering a comprehensive understanding of the network’s properties and dynamics. Specifically, node degree represented the number of neighbors a node has, while node strength accounted for connection weights. The local clustering coefficient quantified interconnectivity among a node’s neighbors. Additionally, the weighted clustering coefficient considered both positive and negative weights [21].

Moving on to node centrality, we utilized four centrality algorithms: degree-based centrality, betweenness centrality [22], PageRank centrality [23], and eigenvector centrality [24]. Degree-based centrality determined central nodes based on the number of connections, while betweenness centrality quantified a node’s position on shortest paths. PageRank and eigenvector centrality considered both connections and neighboring node centrality [25].

Assortativity coefficients were employed to measure the degree of similarity between nodes in terms of their connectivity patterns. These coefficients provided insights into the tendency of nodes to connect with similar or dissimilar nodes based on specific attributes [26]. Positive assortativity coefficients indicated a preference for nodes to connect with similar nodes, while negative coefficients suggested a tendency to connect with nodes possessing different attributes.

Despite the relatively small size of our AVI networks, we also calculated modularity indices to assess the level of segregation or partitioning within the network. Modularity quantified the strength of connections within communities compared to connections between communities [27]. The Louvain algorithm [28] and the Kernighan-Lin fine-tuning algorithm [29] were employed to identify community structures within the AVI network and assess its level of segregation or partitioning.

Additionally, we measured global and nodal efficiency to evaluate information flow and integration within the network, respectively. Global efficiency represented the average inverse shortest paths, while nodal efficiency quantified the efficiency of a node’s neighbors [30]. Mean first passage time [25] provided insights into the general communication efficiency, and search information [31] quantified the energy required for a random walker to find the shortest path between node pairs.

Furthermore, we examined the importance of edges in the network through edge betweenness, which quantified the number of shortest paths passing through a specific edge. By utilizing the Floyd algorithm [32], we computed the shortest path distance and determined the edge betweenness, allowing us to identify critical edges that significantly influenced information flow and communication in the network. This analysis shed light on the network’s structural properties and highlighted key pathways for efficient information transfer.

### 2.5 Feature selection

In addition to conducting descriptive and statistical analyses, our goal was to identify potential features extracted from the AVI network that could predict ADHD in children. To achieve this, we formulated a binary classification task in a supervised learning setting to train predictive machine learning models. Our dataset, denoted as *D* = (*x*_1_*, y*_1_), (*x*_2_*, y*_2_)*, …,* (*x_n_, y_n_*), consisted of labeled data, where each *x_i_* represented a feature vector of dimension *d*, and *y_i_* indicated the binary label (0 or 1) representing whether *x_i_* belonged to control or the ADHD class.

The objective was to learn a function *f* : *R^d^ → {*0, 1*}* that maps each input vector, represented by graph theoretical features *x_i_*, to its corresponding binary label *y_i_*. This function was modeled by a binary classifier that assigned a probability score *P* (*y* = 1*|x*; *θ*) to each input *x_i_*, with *θ* representing the model parameters. To optimize the classifier, our goal was to find the optimal parameters *θ* that either maximized the likelihood of the data *D* or, equivalently, minimized the cross-entropy loss *L*(*θ*) defined as:

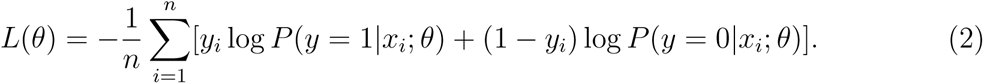

Considering the relatively small number of observations (*n*) compared to the dimensionality of the feature vector (*d*), we implemented two consecutive dimension reduction procedures: analysis of variance (ANOVA) to filter out insignificant features and the minimum redundancy maximum relevance (MRMR) algorithm [33] to identify the most informative features. ANOVA served as a faster approach to filter out insignificant features, while the MRMR algorithm was utilized to identify the most informative features. By considering both the correlation between features and their relevance to the target value, the MRMR algorithm selected a minimal feature set that maximized the overall information content for the model. To ensure better generalizability and prevent data leakage, we adopted a strategy where a held-out test set, comprising 20% of randomly selected observations, was set aside. Furthermore, a distinct seven-fold stratified cross-validation was performed to select relevant features from the training set. Features that consistently demonstrated informativeness across more than two folds were selected for further analyses and for training the final classification models. This approach enhanced the reliability and significance of the selected features for the final model development and training.

Before further processing with final model training, we employed classical statistical methods, including the Shapiro test for normality assessment, independent t-test for normally distributed features, and Mann-Whitney U-test for non-normally distributed features, to examine the chosen features and identify significant distinctions between the ADHD and neurotypical groups. Our aim was to uncover noteworthy global and local measurements, with a *p*-value set at 0.05.

### 2.6 Machine learning classification

Following the feature selection process, we proceeded to train ensemble learning models, which encompassed different variations of random forest and XGBoost. These models were utilized to classify features derived from the AVI networks into ADHD and neurotypical (control) groups. Our objective extended beyond comparing their respective performances; we sought to determine the utmost potential of AVI network alterations as indicators of ADHD in children.

To ensure the generalizability of our results, we reserved the aforementioned 20% of observations as a held-out test set, which remained consistent for all experiments. The remaining data points were utilized for training and validating the models. Given the dataset’s significant class imbalance (ADHD to control ratio of 0.42), we employed stratified cross-validation with 7 folds to ensure that each fold maintains the same class distribution as the original dataset. This approach helps prevent bias and ensures more reliable hyperparameter tuning for the models [34]. The classification performance of the best trained model was ultimately evaluated using the held-out fold test set.

Throughout the process of hyperparameter tuning and model evaluation, we placed greater emphasis on the F1-score rather than accuracy. This approach aimed to mitigate any bias towards the majority class, which in this case is the neurotypical group. In addition to the F1-score, we also report other metrics such as sensitivity, specificity, and the area under the receiver operating characteristic curve (ROC) to assess the models’ ability to accurately predict the presence or absence of ADHD. These measures provide a more comprehensive understanding of the models’ performance and their predictive capabilities.

To perform comprehensive classification of the features extracted from AVI networks into ADHD and neurotypical (control) groups, we employed a diverse set of powerful machine learning models, including balanced random forest (BRF), XGBoost, and EasyEnsemble classifier (EEC).

The random forest algorithm comprises an ensemble of decision trees, where each tree is constructed from a bootstrap sample drawn with replacement from the training set [35]. In addition to the bootstrap sample, a portion of the training data, referred to as the out-of-bag (oob) sample, is reserved for later evaluation. To introduce further variability and reduce correlation among the decision trees, feature bagging is employed, introducing randomness into the dataset to achieve a better robustness. During the prediction process, a majority vote is conducted, wherein the most frequently occurring categorical variable determines the predicted class. The oob sample, which was set aside earlier, plays a crucial role in cross-validation, contributing to the finalization of the prediction.

Random Forest combines the power of bagging as an ensemble method with the utilization of multiple decision trees as weak learners to effectively model the data. However, when dealing with imbalanced datasets, there is a possibility that the bootstrap subsets may contain an excess of data points from the majority class, leading to subpar performance. To address this issue, Chen et al. [36] proposed a balanced random forest algorithm that incorporates resampling techniques such as undersampling within the bootstrap subsets. In our study, we utilized the undersampling version of the balanced random forest.

EasyEnsemble is another ensemble learning technique specifically designed to tackle the difficulties posed by imbalanced datasets [37]. Unlike conventional approaches that simply undersample the majority class, EasyEnsemble classifier takes a more sophisticated approach. It creates multiple subsets from the majority class while retaining all samples from the minority class. These subsets, along with the complete set of minority class samples, are then utilized to train a series of base classifiers.

In the EEC method, adaptive boosting (AdaBoost) is often employed as the underlying algorithm for the base learners [38]. AdaBoost is a sequential learning algorithm that optimizes a linear combination of multiple weak classifiers, each slightly better than random guessing. By leveraging AdaBoost in EEC, the model benefits from the iterative process of adjusting weights and focusing on the challenging samples, thereby enhancing the overall performance of the ensemble.

By utilizing the power of multiple classifiers and AdaBoost’s boosting mechanism, EasyEnsemble classifier effectively addresses the imbalance issue by providing a more nuanced and sophisticated approach to training models on imbalanced datasets. This method aims to capture the intricacies of the minority class while avoiding the pitfalls of under-representing the majority class, resulting in improved classification accuracy and better handling of imbalanced data. In addition to the random forest algorithm and EasyEnsemble technique, we also employed extreme gradient boosting (known as XGBoost) as one of the models for solving the classification task at hand. The XGBoost stands as a highly regarded and remarkable machine learning algorithm renowned for its exceptional performance, efficiency, and scalability. It has garnered popularity and been widely employed in various competitions and real-world applications.

As an ensemble method, XGBoost capitalizes on gradient boosting to achieve its robustness [39]. This technique utilizes different weak learners to model the data, with each weak learner trained to rectify the errors made by preceding models. The essence of the gradient aspect lies in XGBoost’s ability to iteratively train new models by optimizing a differentiable cost function through negative cost gradient descent.

XGBoost offers several advantages, including parallelization, which allows for efficient utilization of computational resources, and regularization, which aids in mitigating overfitting by imposing constraints on model complexity. In this study, we test its effectiveness in handling imbalanced data distribution.

## 3 Results

We employed pre-processed fMRI data that had undergone rigorous preprocessing using the NIAK software [16], ensuring high data quality and minimizing potential artifacts before our analysis.

We began by selecting the average regional time series of 68 ROIs, depicted in Figure 2 from the audiovisual integration network. Connection matrices were constructed using both linear Pearson correlation and the mutual information algorithm. To identify significant patterns or differences between ADHD samples and the control group, we employed the network-based statistics (NBS) test to rigorously examine the functional connectivity profiles.

Density-based thresholding was employed to retain meaningful edges in the reconstructed networks, allowing us to preserve a specified percentage, denoted as *τ*, of the total edges. We fine-tuned the value of *τ* for optimal machine learning classification, as described in the machine learning results section. A wide range of nodal and network-wide graph theoretical measures were applied to the connection matrices, followed by descriptive and statistical analyses to identify potential features for predicting ADHD in children.

Finally, using supervised learning, we formulated a binary classification task to train predictive machine learning models capable of distinguishing between ADHD and neurotypical samples.

The subsequent subsections present the findings of our study.

### 3.1 Network-based statistics

In this study, we aim to explore potential differences in the functional audiovisual integration network patterns between children with ADHD and their neurotypical peers. To do so, we conducted a network-based statistics (NBS) analysis on the functional connectivity data from participants in both groups. The NBS method is advantageous for exploratory investigations as it allows us to work directly with the raw connectivity measures, eliminating the need for thresholding the adjacency matrix.

For robust statistical analysis, we performed a total of 1000 random permutations to estimate the null distribution. The NBS was systematically applied using primary thresholds (t-statistic) ranging from 1 to 15. In all experiments, the identified over-connected and under-connected components, defined based on the ADHD-Control comparison in the NBS analysis, exhibited a highly significant *p*-value of less than 0.001. Notably, increasing the primary thresholds led to the detection of sparser over-connected and under-connected networks. Consequently, for visualization purposes, we optimized the primary threshold parameter to create highly sparse over-connected and under-connected networks, including approximately 1.0% of all possible links (See Figure 3).

**Figure 3:**
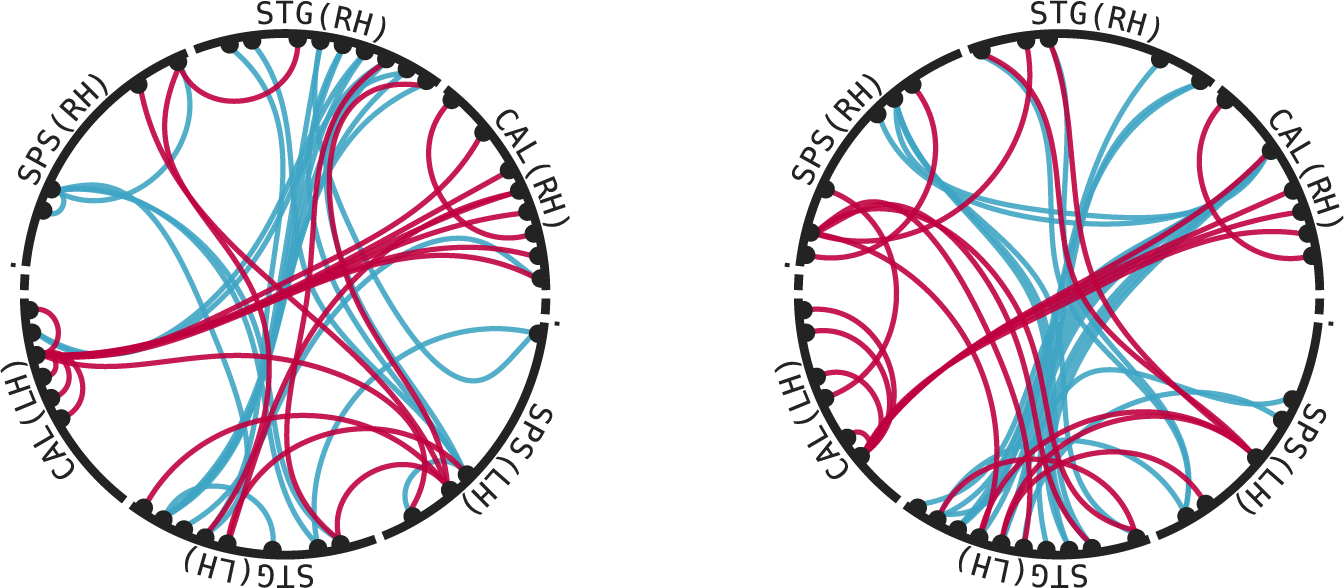
Graphical illustration of significant sub-networks under the over-connectivity (red) and under-connectivity (blue). **Left:**sub-networks identified in MI-driven connectivity networks with *t* = 7 and *t* = 10 for over-connectivity and under-connectivity, respectively. **Right:** sub-networks detected in connectivity profiles obtained from Pearson correlation analysis with *t* = 13 and *t* = 14 for over-connectivity and under-connectivity.

Figure 3 illustrates the identified sub-networks in the mutual information (MI) reconstructed connectivity profiles and the connectivity matrices obtained from computing Pearson correlation coefficients between time series during the entire scanning duration. The NBS analysis was conducted using the Brainconn package [40].

The NBS analysis revealed consistent under-connectivity in edges linking the left superior temporal gyrus (STG) to the superior parietal sulcus (SPS), STG, and calcarine sulcus (CAL) regions in the right hemisphere. Additionally, a few under-connected links were found between STG and SPS regions in the left hemisphere in both network types. In correlation-driven networks, noticeable under-connected links were also detected between the right CAL and SPS regions.

Conversely, for the over-connected component, the sub-networks demonstrated higher connectivity between inter-CAL edges in both hemispheres, as well as edges involving the left STG and left SPS. Furthermore, inter-hemispheric connections were more pronounced in edges connecting the right and left CAL, along with left SPS to right STG. Notably, the connections between the Heschl’s gyrus (HES) region and other nodes are absent from both over- and under-connected sub-networks. As a result, these connections are not depicted in Figure 3.

In MI-constructed connectivity patterns, the over-connected component prominently exhibited connections between the left and right SPS, whereas, in correlation networks, connections between the left STG and right SPS were more evident.

### 3.2 Features statistics

After applying density-based thresholding to the connectivity matrices, a total of 12, 143 graph theoretical features were derived from each connectivity matrix. These measurements were computed on undirected weighted connectivity matrices using the Brainconn package [40].

These variables encompassed five global measurements, 748 node-based measurements, and 11, 390 edge-based measurements. To ensure the selection of robust features, we employed the ANOVA and MRMR algorithms in a separate seven-fold cross-validation process. Initially, the ANOVA algorithm was utilized to select the 1, 000 most promising features. Subsequently, the MRMR algorithm was applied to identify the best features among the selected set. As a final step in feature selection, we retained the features that exhibited importance in more than two folds. This step served to further refine the feature set, resulting in the identification of 40 features in MI-driven connectivity matrices and 35 features in connectivity profiles obtained from Pearson correlation analysis. Selected features are shown in Supplementary Figures S1 and S2.

Prior to training any machine learning classification model, we performed classical statistical analyses on the final set of selected features. Initially, we examined the normality of the data using the Shapiro test. Subsequently, for features that displayed a normal distribution, we utilized an independent t-test. On the other hand, for features that did not exhibit normality, we employed the Mann-Whitney U-test. Our objective was to identify significant global and local measurements, using a significance level of 0.05.

The 40 chosen features from the connectivity matrices created through the mutual information approach primarily fell into three main categories: edge betweenness, number of edges in the shortest path (NESP), and search information. The distribution of these features, along with their respective *p*-values, can be observed in Supplementary Figure S1. Notably, nearly all of these features exhibited significant differences between the ADHD and neurotypical groups. For instance, the mean values of the selected edge betweenness measurements were significantly higher in the ADHD group (*p <* 0.001). On the other hand, the average number of edges in the shortest path (known as characteristic path length) was notably lower in the ADHD group (*p* = 0.006). In terms of the mean search information feature, no significant differences were observed between the two groups. Nonetheless, the search information feature, which measures the energy needed for a random walker to discover the shortest path between pairs of nodes, displayed notable distinctions between the two groups across multiple node pairs (for additional information, refer to Supplementary Figure S1).

Moreover, our feature selection algorithm revealed a multitude of features (17 for MI-driven networks and 16 for correlation-driven networks) capturing the inter-hemispheric connectivity patterns. Among the uni-hemisphere features, those extracted from the right hemisphere displayed slightly greater significance in MI-driven matrices, while in connectivity profiles obtained from Pearson correlation analysis, the features from the left hemisphere were more prominent. These observations align remarkably well with the under-connected and over-connected components detected by the NBS algorithm (Figure 3).

These findings further underscore the significance of the selected features in distinguishing between the ADHD and neurotypical groups. In particular, the significant differences observed in the edge betweenness and NESP measurements suggest significant alterations in resting-state functional connectivity patterns in the audiovisual integration network in children with ADHD compared to their neurotypical peers.

### 3.3 Machine learning results

To further investigate potential differential patterns of associations in audiovisual integration (AVI) network connectivity, we employed a classification task based on extracted features. The objective was to explore whether advanced machine learning models could be used for diagnostic purposes with a limited set of AVI-derived features. Given the dataset’s significant class imbalance (ADHD to control ratio of 0.42), we opted for ensemble learning models, such as random forest, XGBoost, and EasyEnsemble classifier, which were specifically developed to cope with the challenges imposed by an unbalanced number of observations in classes. We implemented our models by leveraging the imbalanced-learn package [41], an extension of the scikit-learn library, which offers a suite of resampling techniques designed to enhance model performance on imbalanced class distributions.

After establishing the connectivity matrices through Pearson correlations and mutual information approaches, density-based thresholding was applied to each connectivity profile to mitigate the impact of spurious links in connectivity matrices. We conducted a grid search and trained all models to find the optimal density threshold. The performance of the models was assessed by calculating the area under the receiver operating characteristic curve (AUC) to ensure the maximum sensitivity and specificity of the model. The density values of 0.90 and 0.87 were found to be the most favorable choices for all models for MI and Pearson-created connectivity matrices, respectively, indicating that preserving approximately 90% of the links effectively captured the relevant connections (see Figure 4).

**Figure 4:**
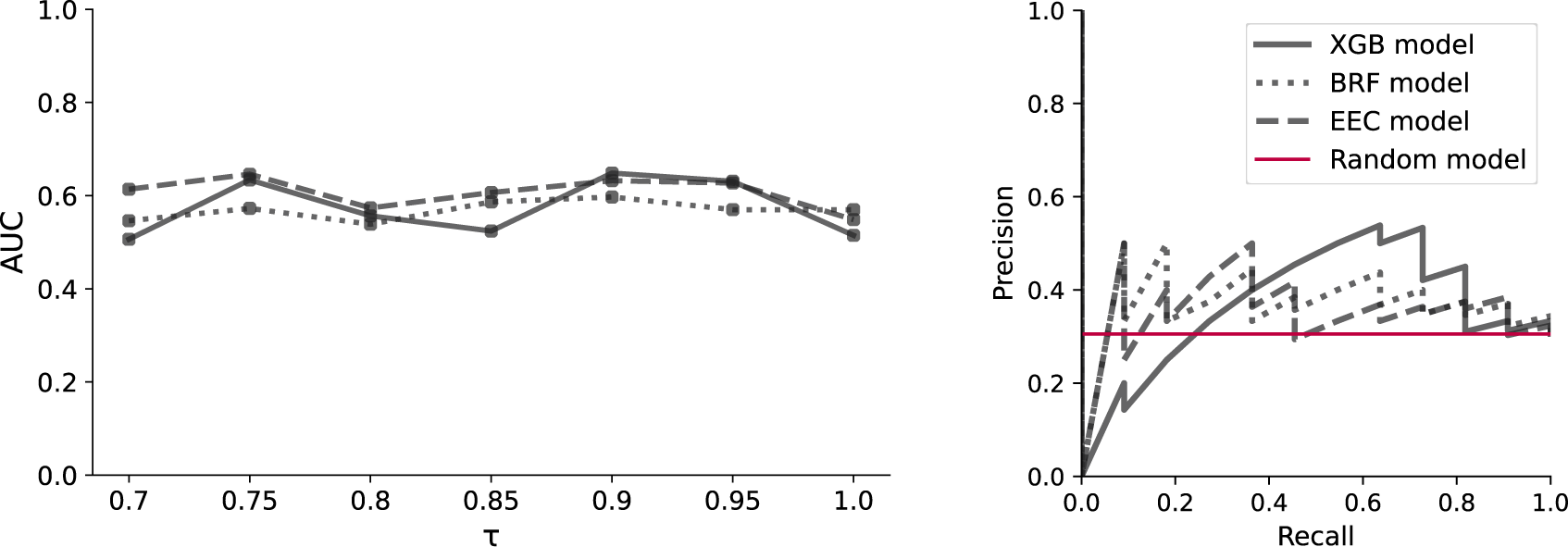
Machine learning models’ performance **Left:** Fine-tuning the percentage of total edges that are preserved during the density-based thresholding process. **Right:** Precision-Recall curves for models trained on the MI-driven AVI networks.

Given the set of selected features for participants in the training set, we optimized hyperparameters of the three ensemble learning classifiers using seven-fold cross-validation and extensive grid search. In all the experiments, 20% of the observations, including 11 ADHD and 25 control samples, were reserved as a held-out test set. The remaining data points were utilized for training and validating the models. All features were normalized by removing the mean and scaling to unit variance before being fed to the ML models.

Each model underwent independent fine-tuning within a predefined search space, utilizing the training data. The overall classification results obtained through cross-validation are presented in Table 1 and 2 (upper) using five performance metrics. These metrics capture various aspects of model performance. In cases of MI, while the EasyEnsemble classifier (EEC) achieved a higher mean AUC compared to the other models, the XGBoost model demonstrated a higher mean sensitivity (i.e., ADHD prediction) and exhibited lower variance in the majority of the evaluation metrics. Our models performed better on the features extracted from resting-state functional connectivity matrices created through the mutual information approach compared to Pearson correlation.

**Table 1:**
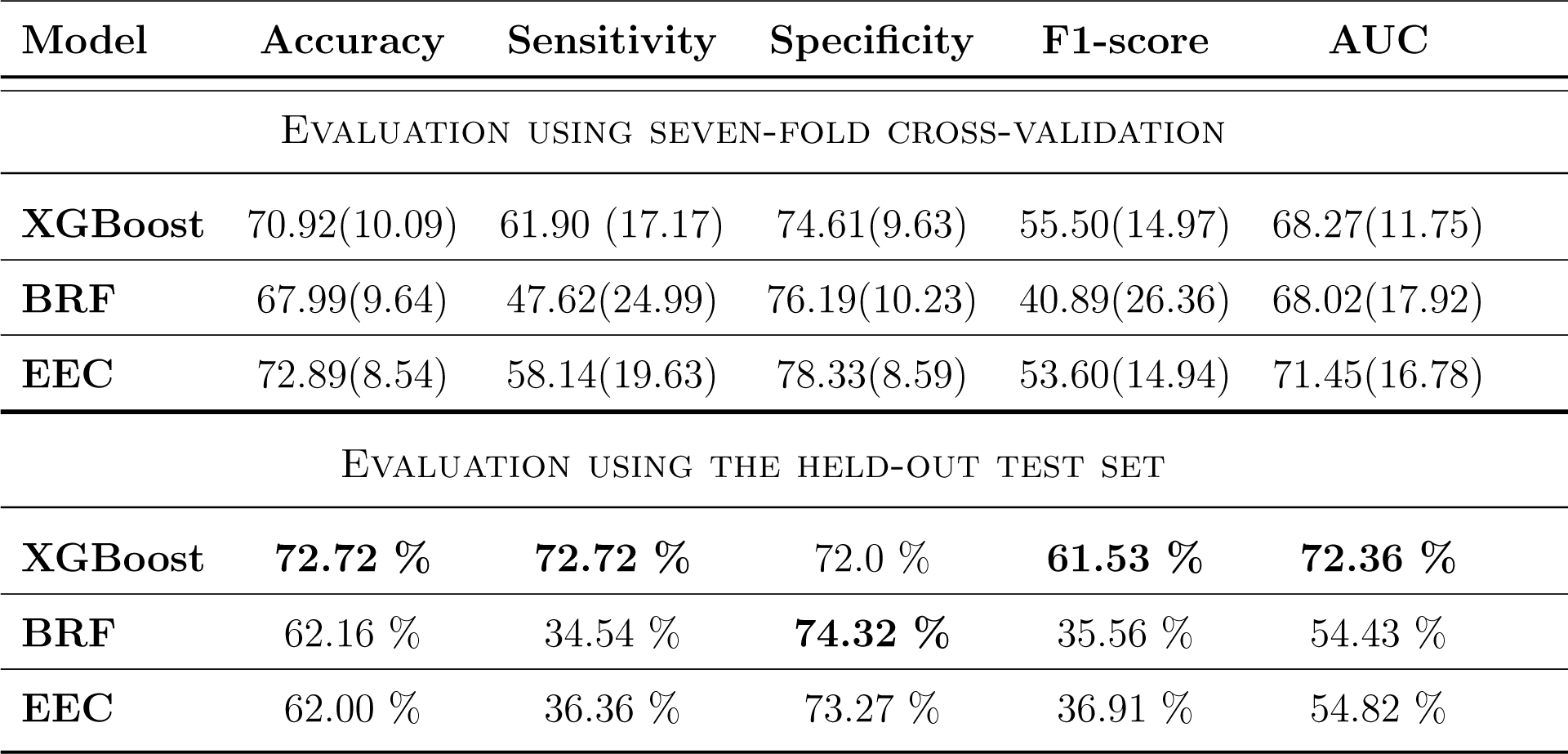
Quantitative evaluation of the classification performance of the considered machine learning models: XGBoost, random forest (BRF) and EasyEnsemble classifier (EEC). Evaluation metrics are: mean class-wise accuracy, ADHD diagnosis sensitivity and specificity, as well as F1-score and AUC to measure the balance between the ability of the model to correctly detect pathological samples and correctly discard control observations. Variance in cross-validation setting is reported in parenthesis. The connectivity matrices are generated through mutual information method.

**Table 2:**
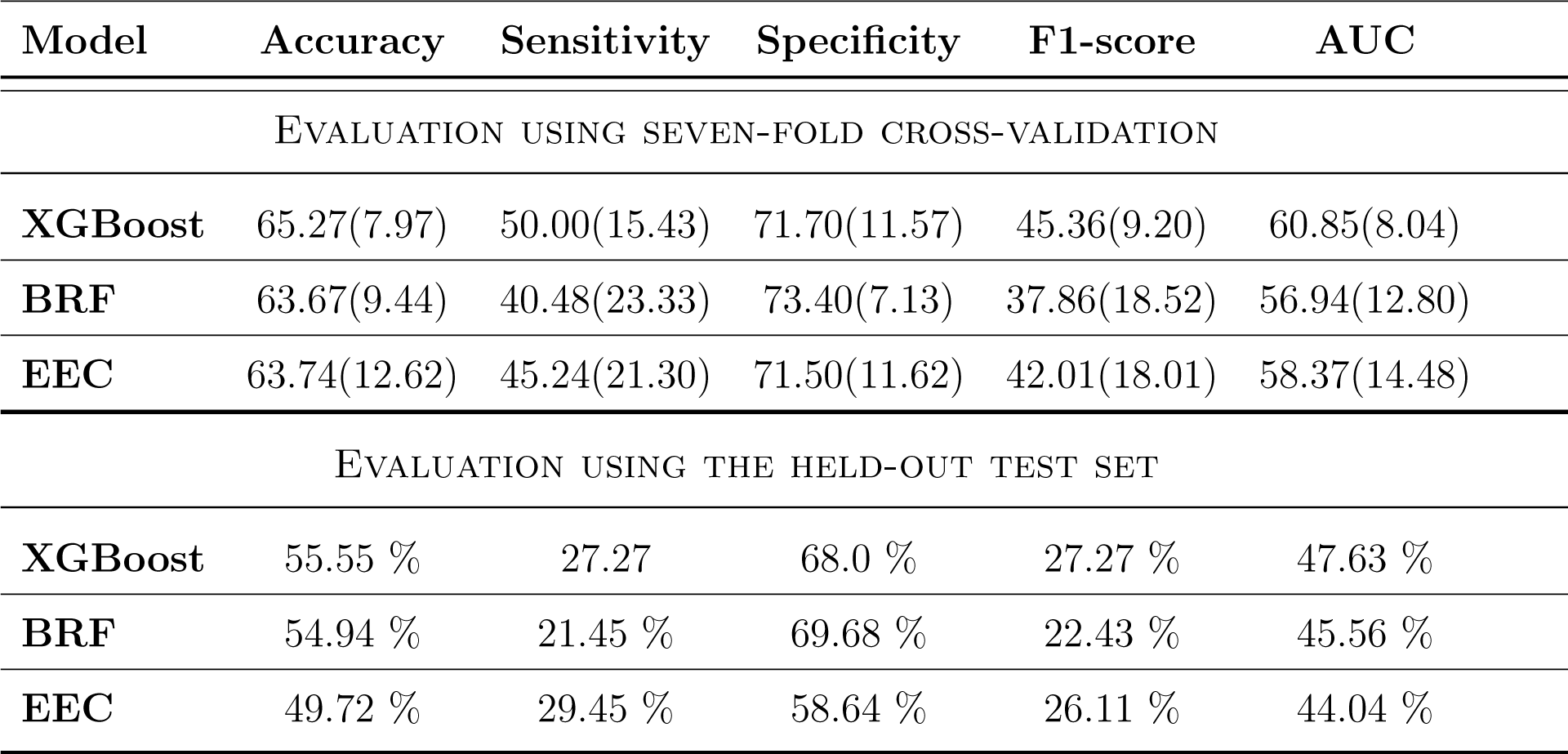
Quantitative evaluation of the classification performance of the considered machine learning models on features extracted from connectivity profiles generated by linear Pearson correlation.

To determine whether the model’s performance exceeded chance levels in this unbalanced set, we constructed precision-recall curves, as depicted in Figure 4, for all the models and both MI and Pearson correlation-created matrices. These graphs corroborate the superiority of XGBoost compared to other models and show its consistency in both MI and Pearson correlation cases.

Finally, we assessed the models’ performance on the held-out test dataset and reported the results in Tables 1 and 2 (lower) for connectivity profiles obtained from MI and Pearson correlation approaches, respectively. In the case of MI (i.e., Table 1 (lower)), the XGBoost model not only exhibits acceptable performance but also ensures a balance between all the evaluation metrics on the test set, confirming that the model did not overfit on the training samples. The other models tend to have higher specificity and low sensitivity despite carefully conducted cross-validation.

Surprisingly, the models perform better when trained on the features extracted from MI connectivity matrices. Moreover, in the case of Pearson correlation, we were not able to find any optimum set of hyperparameters that yields a sensitivity-specificity balance, and hence, higher values of F1-score and AUC.

## 4 Discussion

Previous research on the neural basis of audiovisual integration in the human brain has established a flexible multiple pathways model, highlighting the crucial role of the superior temporal cortex, variations in neural assemblies based on stimulus and attention, and consistent neural activity in this region [42]. However, the distinctive pattern of associations within this network remains unexplored in children with ADHD, despite their known deficits in sensory processing and integration. To bridge this knowledge gap, we conducted a comprehensive study on graph-theory-based analyses of rs-FMRI data to identify potential diagnostic biomarkers for ADHD and gain insights into the connectivity patterns within the audiovisual integration (AVI)-related brain network.

To begin, we selected specific brain regions associated with AVI based on previous literature and their potential involvement in AVI during periods of rest. In our study, we excluded brain regions activated solely during audio-visual sensory tasks, focusing instead on regions potentially engaged in audio-visual integration during resting states. The selected areas demonstrate heightened activity in the simultaneous processing of visual and auditory stimuli, emphasizing intrinsic stimulus features over content attributes. These regions included the superior temporal gyrus/sulcus, superior parietal sulcus, Heschl’s gyrus, and calcarine fissure and its surrounding cortex [8, 9, 10, 11, 13, 12].

Next, we employed a diverse array of measures to comprehensively examine the organization and efficiency of the AVI-related brain network in children with ADHD compared to a neurotypical control group. It is important to note that our approach was primarily exploratory, rather than driven by specific hypotheses. Consequently, it is natural that we observed variations in specific metrics, including edge betweenness, number of edges in the shortest path (NESP), and search information, when comparing the two groups.

Given the exploratory nature of our study and the inherent challenge of statistically controlling for these variations, it is essential to acknowledge these observed differences. Furthermore, it is worth highlighting that these metrics play a crucial role in our subsequent analysis, where they are used as features to train machine learning models for classifying subjects into control and ADHD groups. While we acknowledge that some of the observed differences may be influenced by random variability, these variations can still offer valuable insights into potential distinct patterns of AVI-related brain connectivity in children with ADHD.

To further validate and refine our findings, we recommend future research endeavors to conduct confirmatory studies with predefined hypotheses and the use of appropriate correction methods for multiple comparisons. These studies can build upon our exploratory findings and provide a more robust basis for understanding the underlying brain network differences between the two groups.

Alongside the extensive graph theoretical exploration, our network-based statistics (NBS) analyses focused on the raw connectivity matrices to identify specific connections that exhibited significant differences between the two groups. Our findings emphasize the distinctive connectivity patterns associated with both under-connectivity and overconnectivity in the audiovisual integration network, regardless of the method used to construct the connectivity matrices. Furthermore, we posit that these observed discrepancies may elucidate the basis for the distinct performance disparities exhibited by individuals with ADHD in tasks pertaining to audiovisual integration when contrasted with neurotypical control cohorts.

In particular, the discerned state of over-connectivity with the Calcarine (CAL) region prompts the inference of a pronounced visual stimulus bias among individuals with ADHD. This supposition finds concurrence with previous research by Panagiotidi et al. [43], which proposes a tendency for individuals manifesting heightened ADHD-like traits to demonstrate a predilection for visual stimuli, particularly under visuoauditory conditions.

Moreover, our analyses reveal that connections attributed to the left superior temporal gyrus (STG) persist across both under-connected and over-connected network components. Considering the shared occupancy of this region by the audiovisual integration network and that implicated in language-centric functions, the identified perturbations hold the potential to elucidate the deficits observed within language-related functionalities in the context of audiovisual conditions among individuals with ADHD. Notably, a corollary is drawn from existing literature highlighting that children historically afflicted with specific language impairment evince exacerbated performance deterioration in simultaneous audiovisual stimulus detection when the ADHD index registers higher [44]. Additionally, earlier research has demonstrated that the left hemisphere becomes notably engaged during lipreading, as an example of audiovisual speech perception [45]. Based on this knowledge and considering the trajectory of audiovisual speech perception in the early stages of development, we propose that the variations in connectivity patterns within the left STG region could potentially provide insight into the challenges observed in AVI tasks related to language among individuals with ADHD.

To go beyond descriptive and statistical analyses, we applied machine learning techniques to differentiate between the control group and children with ADHD based on the extracted graph theoretical features served as inputs. We evaluated the performance of multiple ensemble learning models through cross-validation, which involved splitting the data into training and validation sets and iteratively training the models on different subsets of the data. Additionally, a held-out test set was used to assess the generalization ability of the selected model.

Notably, all the classifiers trained on the AVI network drawn from the mutual information approach outperformed classifiers trained on the connectivity patterns derived from the linear Pearson correlation method. Furthermore, comparing the cross-validation and test results, the classifiers trained on the AVI network drawn from the mutual information approach seem to be more consistent and less prone to overfitting. We speculate that the ability of mutual information to capture nonlinear relationships or its robustness to noise in the data may explain this superiority. During cross-validation, all the models exhibited a notable degree of variability in their performance, which can be attributed to two key factors. Firstly, the heterogeneity among samples within each class, as outlined by Zhou et al [46] in their study, poses a significant challenge. This means that samples within the same class, such as individuals with ADHD or the control group, displayed considerable variations in characteristics or features that the models found challenging to account for. Secondly, the imbalanced distribution of samples across different classes within each crossvalidation fold further contributed to this variance. Moreover, it’s important to recognize that this variance appears less pronounced when assessing models solely based on their classification accuracy. To draw more robust and comprehensive conclusions about model performance, it is essential to take into consideration multiple evaluation metrics beyond just accuracy, including metrics like sensitivity, specificity, and precision, as they provide a more nuanced perspective on model effectiveness, especially when dealing with imbalanced datasets.

A recent review of diagnostic ML models developed for ADHD using the same rs-fMRI data in the ADHD-200 dataset shows that in the classification setting with a held-out test set, the accuracy of the models ranges from 37% to 73% when the regions of interest (ROIs) are selected from the whole brain network [47]. In this context, our study suggests a pre-processing and analysis pipeline that achieves a comparable performance by only selecting the ROIs in the AVI network. Unfortunately, in these studies, the more relevant metrics such as sensitivity and F1-score are not commonly reported. One study that reports sensitivity based on the data collected from the center that is used in our study reports a maximum sensitivity of 55.6% while specificity and accuracy are reported as 83% and 59%, respectively [48]. This study introduces a deep Bayesian network in which the deep belief network (DBN) is applied to normalize and reduce the dimension of the rs-fMRI data in every Brodmann area, and the Bayesian network is used to extract the feature of relationships in the whole brain network by structure learning.

To put our results into a better perspective, we also conducted a similar analysis on the default mode network (DMN), which is considered the de facto standard in rs-fMRI studies. Details of this implementation are introduced in Supplementary Note 1. Remarkably, the classifiers trained on the AVI network outperformed classifiers trained on the default mode network. This finding suggests that AVI-related features have superior potential in capturing distinctive patterns of ADHD. By employing a rigorous evaluation approach and demonstrating the superior performance of the classifiers on the AVI network, we aimed to obtain reliable and unbiased estimates of classification performance and highlight the diagnostic potential of AVI-related features.

In our study, we introduced a method that addresses a binary classification task based on graph theoretical features, leveraging ensemble learning models. An interesting aspect of our approach is its potential for extension to ordinal multi-class classification tasks, provided that labeled data are available. This adaptability underscores the versatility of our methodology for addressing various classification challenges beyond the scope of this study. In this regard, an intriguing aspect related to our study is the complex nature of ADHD, often considered a spectrum disorder. This prompts consideration of how our findings might be influenced by shifting diagnostic thresholds or viewing ADHD as a continuous spectrum. These questions underline the need for further exploration into the relationship between our observed network differences and the clinical spectrum of ADHD, providing a path for future investigations into the condition’s nuances.

## 5 Conclusion

In this study, we developed a comprehensive processing pipeline to investigate the intricacies of the audiovisual integration (AVI) network within the context of ADHD in children. This innovative approach has provided valuable insights and has enabled the development of machine learning (ML) diagnostic classifiers that outperform traditional models based on the default mode network, commonly used in children’s ADHD diagnostic studies.

A key aspect of our investigation was to explore the significance of measuring interactions between regional signal time-series using both the non-linear mutual information approach and the linear Pearson correlation method in the study of rs-fMRI-based ADHD diagnostic classifiers. By leveraging these complementary techniques, we sought to better understand the complex dynamics of the AVI network and its role in ADHD pathology.

The classification accuracies obtained in our study align well with previous research on whole brain network analysis [48]. Additionally, we adopted ensemble learning models designed to address challenges posed by unbalanced samples, fine-tuning hyperparameters to achieve an optimal balance between sensitivity and specificity. However, it is worth noting that despite these efforts, the overall classification performance remains moderate, likely due to the inherent neurobiological heterogeneity within the audiovisual integration network and its correlation with developmental stages. Moreover, the limited sample size and imbalanced data continue to pose significant obstacles in developing more accurate and clinically useful imaging classifiers for ADHD.

Looking ahead, we propose that future classification studies utilizing a similar pipeline should explore multimodal data integration, combining functional, structural, and/or diffusion tensor imaging MRI. This approach promises to enhance our comprehension of AVI network alterations in ADHD and address the phenotypic heterogeneity commonly observed in clinical populations.

Taken together, our findings provide compelling evidence of aberrations within the audiovisual integration network in children with ADHD. The identification of AVI network partial hypo- and hyper-connectivity underscores the critical significance of incorporating these observations into neural models of ADHD, thereby advancing our understanding of the disorder and its underlying mechanisms.

## 6 Acknowledgments

Fatemeh Hadaeghi’s work was supported by the German Research Foundation (DFG) TRR169-A2. The authors affiliated with the University of Shahid Beheshti were funded by the Cognitive Sciences and Technologies Council (Grant Number: 9875). We acknowledge Claus Hilgetag’s significant role in our project, which included engaging discussions and valuable feedback on the manuscript, contributing significantly to the strength of our research.

## 8 Author contributions

Conceptualization: MZ, FH, AP, FB. Data curation: MZ. Formal analysis: MZ, FH. Investigation: MZ, AP, FH. Methodology: MZ, FH. Supervision: FH, FB. Validation: AP, FH. Visualization: MZ. Writing – original draft: FH, MZ. Writing – review & editing: MZ, AP, FB, FH.

## 9 Data and materials availability

The code necessary to replicate the experiments and all pertinent data will be made available upon acceptance of the manuscript or upon request by the reviewers. Please contact the corresponding author for access to the GitHub repository containing the code and data.

