## Supplemental Document: Note S1, Tables S1, S2, and S3, Figs. S1 and S2 for "Differential Patterns of Associations within Audiovisual Integration Networks in Children with ADHD"

MOHAMMAD ZAMANZADEH[1], ABBAS POURHEDAYAT[1], FATEMEH BAKOUIE[1], AND FATEMEH HADAEGHI[2,*]

[1] *Institute for Cognitive and Brain Sciences, Shahid Beheshti University, Tehran, Iran*
[2] *Department of Computational Neuroscience, University Medical Center Hamburg-Eppendorf, Hamburg, Germany*

### 1. SUPPLEMENTARY NOTE 1: STUDY ON DEFAULT MODE NETWORK (DMN)

The primary hypothesis of this study proposes that there are distinct differences in the developmental pattern of the functional audiovisual integration (AVI) network between children with Attention Deficit Hyperactivity Disorder (ADHD) and their neurotypical peers. Our analysis, using network-based statistics (NBS) and machine learning classification, confirms this alteration and demonstrates that features extracted from the AVI network have some degree of predictive ability for ADHD. To put our machine learning results into a better perspective, we conducted similar analyses on the default mode network (DMN), which is considered the standard in ADHD-related resting-state functional MRI (rs-fMRI) studies [1].

To reconstruct the resting-state functional connectivity of the DMN, for the sake of consistency, we carefully selected 68 regions of interest (ROIs) distributed across six brain regions in both hemispheres. Specifically, we included the posterior cingulate cortex (PCC), precuneus (PCUN), medial prefrontal cortex (MPFC), inferior parietal lobule (IPG), middle temporal gyrus (MTG), and inferior temporal gyrus (ITG).

Following the steps of feature extraction and selection in our processing pipeline, we optimized and trained our machine-learning classification models. Subsequently, we evaluated the model with the best performance through cross-validation on the held-out test set. The evaluation metrics are summarized in Table S1.

**Table S1.** Quantitative evaluation of machine learning models' classification performance based on features extracted from the MI reconstructed connectivity profile of the Default Mode Network (DMN).

| Model | Accuracy | Sensitivity | Specificity | F1-score | AUC |
|---|---|---|---|---|---|
| EVALUATION USING SEVEN-FOLD CROSS-VALIDATION | | | | | |
| **XGBoost** | 66.73(14.18) | 52.38 (22.59) | 72.79(18.04) | 48.09(18.30) | 62.59(14.24) |
| **BRF** | 66.09(13.07) | 42.86(12.14) | 75.71(18.72) | 43.20(13.21) | 59.29(10.17) |
| **EEC** | 64.52(12.76) | 52.38(25.86) | 69.66(12.92) | 45.39(18.90) | 61.02(15.33) |
| EVALUATION USING THE HELD-OUT TEST SET | | | | | |
| **XGBoost** | 63.88 % | 68.0 % | 54.54 % | 57.99 % | 61.27 % |

**Table S2.** The precise coordinates $(x, y, z)$ of 68 regions of interest (ROIs) chosen from both hemispheres to examine the audio-visual integration (AVI) network.

| Region | Coordinate (RH) | Coordinate (LH) |
|---|---|---|
| CAL 1 | (14.68, -63.99, 10.42) | (-10.28, -75.87, 11.09) |
| CAL 2 | (5.56, -81.03, 10.42) | (-7.16, -73.41, 11.09) |
| CAL 3 | (14.68, -65.13, 17.05) | (-13.05, -82.34, 5.46) |
| CAL 4 | (16.01, -88.16, 9.76) | (-9.14, -75.87, 14.73) |
| CAL 5 | (7.94, -81.8, 10.42) | (-9.48, -64.14, 11.09) |
| CAL 6 | (23.62, -65.13, 4.47) | (-12.12, -75.87, 7.11) |
| CAL 7 | (16.26, -98.86, -1.15) | (-5.83, -90.14, -4.79) |
| CAL 8 | (16.26, -98.86, -1.15) | (-9.88, -68.34, 11.75) |
| CAL 9 | (8.34, -90.14, 4.47) | (-22.39, -61.49, 10.09) |
| CAL 10 | (5.74, -69.93, 10.42) | (-4.51, -85.78, -1.82) |
| SPS 1 | (17.87, -60.27, 64.24) | (-20.1, -55.16, 63.59) |
| SPS 2 | (39.65, -52.35, 64.59) | (-22.86, -60.27, 50.4) |
| SPS 3 | (21.37, 58.86, 69.59) | (-18.38, -64.89, 68.53) |
| SPS 4 | (21.83, -60.27, 62.93) | (-30.58, -68.51, 54.35) |
| SPS 5 | (13.01 -81.16 50.70) | (-20.36, -68.51, 48.42) |
| SPS 5 | (13.01 -81.16 50.70) | (-20.36, -68.51, 48.42) |
| SPS 6 | (18.99, -64.64, 52.38) | (-19.37, -68.51, 44.79) |
| SPS 7 | (37, -48.07, 56) | (-24.65, -68.51, 53.03) |
| SPS 8 | (31.23, -67.8, 56) | (-37.87, -44.89, 58.97) |
| SPS 9 | (26.5, -65.88, 52.38) | (-26.63, -47.41, 65.56) |
| SPS 10 | (40.32, -51.21, 56) | (-26.63, -55.16, 48.42) |
| SPS 11 | (29.41, -65.43, 59.63) | (-28.94, -60.27, 65.23) |
| SPS 12 | (15.04, -49.39, 63.59) | (-20.36, -47.41, 63.59) |
| HES 1 | (45.19, -19.60, 9.43) | (-44.24, -22.42, 10.42) |
| STG 1 | (64.62, -20.40, -0.50) | (-57.84, -48.93, 16.06) |
| STG 2 | (54.31, -20.40, -4.14) | (-50.70, -21.98, 8.44) |
| STG 3 | (60.38, -1.57, -11.09) | (-48.33, -30.70, 8.44) |
| STG 4 | (56.07, -45.27, 13.74) | (-60.46, -26.74, 15.40) |
| STG 5 | (45.15, -34.27, 7.45) | (-57.48, -13.27, 2.15) |
| STG 6 | (63.03, -7.86, -4.14) | (-65.76, -47.26, 16.06) |
| STG 7 | (66.60, -44.61, 19.04) | (-59.03, -18.42, 2.15) |
| STG 8 | (67.79, -31.10, 7.45) | (-55.46, -0.24, -10.76) |
| STG 9 | (43.61, -15.64, -0.50) | (-55.46, 1.74, -3.14) |
| STG 10 | (49.95, -7.72, -0.50) | (-61.40, -8.91, 2.15) |
| STG 11 | (51.14, -30.31, 12.42) | (-68.14, -16.83, 2.15) |

**Table S3.** A diverse range of local and global graph theoretical measures were conducted on the reconstructed connection matrices. These measures encompassed three main categories, including global features, nodal variables, and edge-based attributes. LCAV: Louvain refined community affiliation vector, KCAV: Kernighan-Lin community affiliation vector, WSPL: weighted shortest path length, SP: shortest path, MFPT: first passage time, NESP: number of edges in the shortest path.

| | Feature name | Number of features | Reference |
|---|---|---|---|
| **Global features** | Assortativity | 1 | [2] |
| | Louvain modularity | 1 | [3] |
| | Global efficiency | 1 | [4] |
| | Modified modularity | 1 | [5] |
| | Density | 1 | |
| **Node-based features** | Clustering N (negativity weights) | 68 | [6] |
| | Clustering P (positive weights) | 68 | [6] |
| | Clustering coefficient | 68 | [7] |
| | Betweenness centrality | 68 | [8] |
| | Eigenvector centrality | 68 | [9] |
| | Pagerank centrality | 68 | [9] |
| | Degree strength | 68 | |
| | Degree number | 68 | |
| | Local efficiency | 68 | [4] |
| | LCAV | 68 | [5] |
| | KCAV | 68 | [3] |
| **Edge-based features** | Edge Betweenness | 2278 | [9] |
| | Search Information | 2278 | [10] |
| | WSPL | 2278 | [11] |
| | MFPT | 2278 | [9] |
| | NESP | 2278 | [11] |

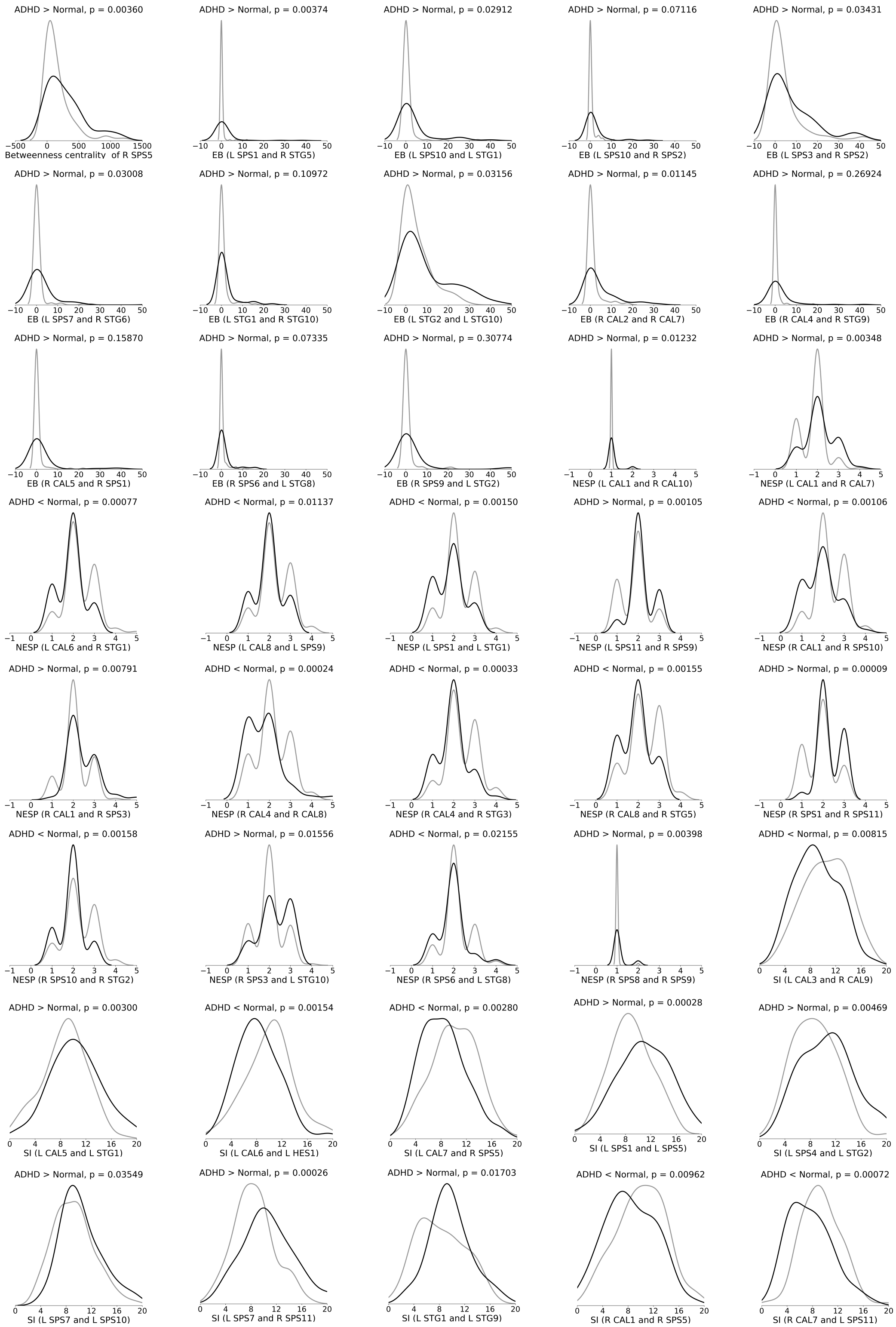


**Fig. S1.** Distribution of 40 features in ADHD (black) and control (gray) groups selected for training machine learning models. Connectivity profiles were created through mutual information. LCAV: Louvain refined community affiliation vector, KCAV: Kernighan-Lin community affiliation vector, WSPL: weighted shortest path length, SP: shortest path, MFPT: first passage time, NESP: number of edges in the shortest path, EB: edge betweenness, and SI: search information

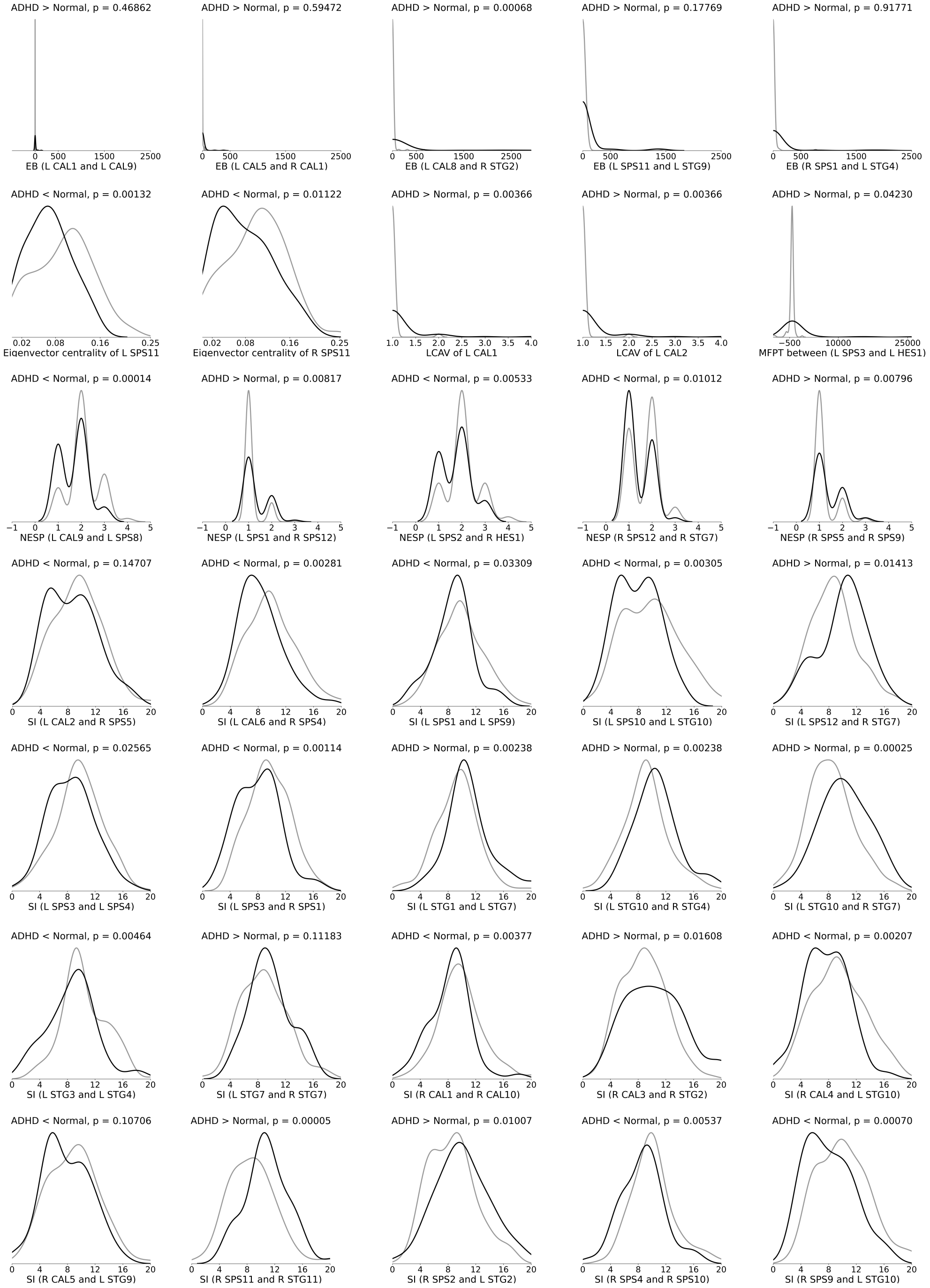


**Fig. S2.** Distribution of 35 features in ADHD (black) and control (gray) groups selected for training machine learning models. Connectivity profiles were created through Pearson correlation. LCAV: Louvain refined community affiliation vector, KCAV: Kernighan-Lin community affiliation vector, WSPL: weighted shortest path length, SP: shortest path, MFPT: first passage time, NESP: number of edges in the shortest path, EB: edge betweenness, and SI: search information
